# Functional brain connectivity during social attention predicts individual differences in social skill

**DOI:** 10.1101/2023.05.19.541349

**Authors:** Samantha R. Brindley, Amalia M. Skyberg, Andrew J. Graves, Jessica J. Connelly, Meghan H. Puglia, James P. Morris

**Author notes:** Corresponding author: James P. Morris, Corresponding author. Authors contributed equally (Meghan H. Puglia, James P. Morris).

## Abstract

Social attention involves selectively attending to and encoding socially relevant information. We investigated the neural systems underlying the wide range of variability in both social attention ability and social experience in a neurotypical sample. Participants performed a selective social attention task while undergoing fMRI and completed self-report measures of social functioning. Using connectome-based predictive modeling, we demonstrated that individual differences in whole-brain functional connectivity patterns during selective attention to faces predicted task performance. Individuals with more cerebellar-occipital connectivity performed better on the social attention task, suggesting more efficient social information processing. Then, we estimated latent communities of autistic and socially anxious traits using exploratory graph analysis to decompose heterogeneity in social functioning between individuals. Connectivity strength within the identified social attention network was associated with social skills, such that more temporal-parietal connectivity predicted fewer challenges with social communication and interaction. These findings demonstrate that individual differences in functional connectivity strength during a selective social attention task are related to varying levels of self-reported social skill.

## Introduction

Social attention is the selection and encoding of social cues (Happé, Cook & Bird, 2017), which is crucial for social cognitive processes including emotion recognition and theory of mind (Puce & Bertenthal, 2015). Consequently, social cues are a highly salient class of stimuli (Wang & Adolphs, 2017) such that human infants preferentially attend to faces and face-like stimuli over inverted faces and face-like stimuli within the first week of life (Farroni et al., 2005).

However, the extent to which social cues automatically capture attention is highly variable in the general population, leading to a range of social attention abilities. For example, autistic individuals often show reduced attention to social stimuli, display deficits in emotion recognition and theory of mind, and experience difficulties during social interactions (Chita-Tegmark, 2016; Hobson, 1986; Baron-Cohen, Alan & Frith, 1985).

Prior research has investigated the neural mechanisms underlying social attention by examining differences in brain activation between autistic and neurotypical volunteers. For example, Herrington and colleagues compared autistics and neurotypicals on a one-back, selective social attention task in which participants viewed composite images of faces overlaid upon houses (Herrington et al., 2015). When attending to faces, autistic participants had increased brain activity in multiple prefrontal regions, including dorsolateral prefrontal cortex (DLPFC). Since DLPFC is a key node in the attentional control network (Hopfinger et al., 2000), increased activity may serve as a potential compensatory mechanism that allows autistic individuals to overcome diminished perceptual representation of faces. Despite group differences between autistics and neurotypicals, DLPFC activity was not predictive of behavioral performance on the social attention task. This highlights a frustrating limitation of traditional fMRI approaches in clinical neuroscience, whereby activity in candidate brain regions reflects group processing differences but fails to capture individual variability associated with behavior.

Brain-behavior associations may be strengthened by also considering functional connectivity, which is frequently used to identify brain networks that support a broad range of complex behaviors. Employing a similar selective social attention paradigm, our group found an intriguing relationship between regional DLPFC activation, salience network functional connectivity, and task performance (Puglia et al., 2018). Specifically, participants that had low functional connectivity within the salience network showed improved performance on the social attention task with more DLPFC activation. In contrast, participants that had high functional connectivity within the salience network showed worse performance with more DLPFC activation. Behavioral performance was not only related to the degree of activation in a specific region of interest, but also to functional connectivity between brain regions.

Individual differences in patterns of functional connectivity can be used to predict behavior, as patterns of functional connectivity are unique to individuals and stable across time (Finn et al., 2015). Connectome-based predictive modeling (CPM) is one example of a cross-validated machine learning approach that assesses individual variability in whole-brain functional connectivity patterns to predict cognitive abilities and traits (Shen et al., 2017). For example, Rosenberg and colleagues utilized functional connectivity during a sustained attention task to predict behavioral performance and applied the results of that CPM model to predict symptoms of attention deficit hyperactivity disorder (Rosenberg et al., 2016). This approach employed a sustained attention fMRI task that amplified individual differences in functional connectivity patterns predictive of stable, related traits (Greene et al., 2018).

Functional connectivity models trained to predict cognitive abilities using task performance are often more robust than models trained to predict traits using self-report questionnaires (Marek et al., 2022). The selection of behavioral assessments with demonstrated reliability and sensitivity to individual differences is essential for generating successful models of out-of-scanner traits (Rosenberg & Finn, 2022). One approach for further decomposing behavioral heterogeneity between individuals is to apply a network psychometrics method to reduce questionnaire data dimensionality and detect distinct communities of traits within a complex phenotype (Lombardo, Lai & Baron-Cohen, 2019; Christensen & Golino, 2021a). Task-induced brain state manipulation and rigorous behavioral phenotyping are complementary efforts to improve brain-based model predictions of real-world traits.

In the present study, we first tested the hypothesis that task-based functional connectivity would reliably predict behavioral performance on a selective social attention task. While we hypothesized that functional connectivity with brain regions previously implicated in social attention, such as the prefrontal cortex, would emerge as important, we also anticipated that successful social attentional mechanisms would require coordinated activity beyond traditionally involved brain regions and networks. Next, we tested the hypothesis that individual differences in functional connectivity patterns during social attention would reliably predict subclinical autistic and socially anxious traits. To test these hypotheses, neurotypical individuals performed an fMRI social attention task (Puglia et al., 2018) and completed questionnaires to quantify the occurrence of autistic and socially anxious traits.

## Methods

### Participants

Sixty-seven individuals (25 males) aged 18-31 (*M* = 22.52, *SD* = 3.70) performed a social attention task during fMRI data acquisition. All participants self-reported as Caucasian American. Recent evidence from machine learning highlights the failure of brain-behavior prediction models to generalize across race (Li et al., 2022). All participants provided written informed consent for a protocol approved by the University of Virginia Institutional Review Board for Health Sciences Research and were paid $50.

### Social attention task

All participants completed a selective social attention task while undergoing fMRI (Puglia et al., 2018). In this one-back matching task, participants either selectively attend to faces or selectively attend to houses in composite images of faces and houses while making a same or different judgment about the previous image. At the beginning of each block, participants saw a prompt instructing them to either selectively attend to faces (Attend Faces condition) or selectively attend to houses (Attend Houses condition) in the composite images. Participants completed six blocks each of the Attend Faces and Attend Houses conditions. Each block consisted of 10 images with either four or five same hits and lasted 40 seconds. The order and pairing of face and house images were randomized for each participant. Block order was pseudorandomized for each participant such that blocks always alternated between Attend Faces and Attend Houses conditions. Stimuli were presented for 1800 milliseconds with an inter-stimulus interval ranging from 200-2400 milliseconds during which a white crosshair was displayed on a black background. Participants responded same or different via button press while the image was still on the screen. Before completing this task, participants completed a practice one-back matching task to ensure they understood the instructions.

Performance on the social attention task was assessed with sensitivity (d’) using the formula: *d’ = z(hit rate) - z(false alarm rate)*. A hit occurred when participants responded same and the correct response was same, while a false alarm occurred when participants responded same but the correct response was different. Due to extreme hit and false alarm proportions, correction prior to d’ calculation was necessary. The log linear approach was applied to prevent z-scores from taking on infinite values by adding 0.5 to both the number of hits and the number of false alarms and adding 1 to both the number of signal trials and the number of noise trials (Hautus, 1995). D’ was calculated separately for the Attend Faces condition and the Attend Houses condition. In both instances, higher d’ values indicated better performance on the social attention task.

### Autistic and socially anxious traits

To quantify the occurrence of autistic and socially anxious traits, an independent sample of 1,357 participants completed five self-report questionnaires. Participants included 593 males and 749 females aged 16-30 (*M* = 19.37, *SD* = 1.69) of various self-reported races (*N* = 1,081 White, *N* = 122 Asian or Pacific Islander, *N* = 50 Black, *N* = 27 Hispanic or Latino, *N* = 3 Native American or American Indian, and *N* = 36 other). Sex was not reported for 15 individuals, and age and race were not reported for 38 individuals. All participants provided written informed consent for a protocol approved by the University of Virginia Institutional Review Board for Social and Behavioral Sciences and received $5 or partial course credit.

The Autism-Spectrum Quotient Questionnaire (AQ) consists of five 10-item subscales that each measure a distinct aspect of the autistic phenotype (Baron-Cohen et al., 2001). We included the social skills, attention switching, communication, and imagination AQ subscales in our analyses. Higher scores on AQ subscales indicated more autistic-like traits and behaviors, specifically fewer social skills, less attention switching behavior and a stronger focus of attention, fewer communication skills, and less imagination, respectively. The Broader Autism Phenotype Questionnaire (BAPQ) consists of three 12-item subscales that each measure traits reflective of genetic liability to autism (Hurley, et al., 2007). We included the aloof personality, rigid personality, and pragmatic language BAPQ subscales in our analyses. Higher scores on BAPQ subscales indicated more autistic-like traits and behaviors, specifically less interest or enjoyment in social interaction, more difficulty adjusting to change, and less effective communication with others, respectively.

The Brief Fear of Negative Evaluation Scale (BFNE) is a 12-item measure that assesses apprehension at the prospect of being treated negatively (Leary, 1983), the Social Interaction Anxiety Scale (SIAS) is a 20-item measure that assesses anxiety experienced during social interactions (Mattick & Clarke, 1998), and the Social Phobia Scale (SPS) is a 20-item measure that assesses fear of being observed or scrutinized by others (Mattick & Clarke, 1998). Higher scores on the BFNE, SIAS, and SPS indicated more social anxiety and phobia, and we included total scores on these questionnaires in our analyses.

### Bootstrap exploratory graph analysis

Bootstrap exploratory graph analysis (bootEGA) using the *EGAnet* package in R (Golino & Christensen, 2021) was employed to reduce the dimensionality of autistic and socially anxious trait questionnaire data into latent space for the 1,357 participants in the independent sample (Christensen & Golino, 2021a). The parametric bootstrap procedure began with exploratory graph analysis (EGA), a community detection and network analysis method that estimates and evaluates dimensional structure (Golino & Epskamp, 2017). The EGA model used the graphical LASSO (GLASSO) procedure to estimate a sparse regularized partial correlation matrix of the collection of variables (Friedman, Hastie & Tibshirani, 2008) where pairwise deletion was implemented to handle missing data (AQ: *N* = 17, BAPQ: *N* = 209, BFNE: *N* = 502, SIAS: *N* = 300, SPS: *N* = 503). Then, bootEGA generated replicate data from a multivariate normal distribution with the same number of cases as the original data, and EGA was applied to each replicate sample for 500 iterations. From this sampling distribution of EGA networks, a typical network structure was estimated by computing the median value of each edge across the replicate networks. Next, the Louvain community detection algorithm was applied to the typical EGA network to define community membership (for comparison, identical community membership was defined by the Walktrap algorithm; Pons & Latapy, 2006). Item stability was estimated by computing the proportion of times each item was placed in each community, which is a metric of structural consistency.

For participants in the current sample with both neuroimaging and questionnaire data (*N* = 65), we computed standardized network scores for each community derived from the EGA model-implied graph. The network scores were computed as a linear combination of the weighted items from the original data that loaded onto each community. Standardized community network scores from EGA are analogous to component vectors from principal components analysis or factor scores from factor analysis (Christensen & Golino, 2021b).

### Imaging parameters and preprocessing

MRI scanning was performed at the University of Virginia Fontaine Research Park on a Siemens 3 Tesla MAGNETOM Prisma Fit high-speed imaging device equipped with a 32-channel head-coil. First, high-resolution T1-weighted anatomical images were acquired using Siemens’ magnetization-prepared rapid-acquired gradient echo (MPRAGE) pulse sequence with the following specifications: echo time (TE) = 2.98 ms; repetition time (TR) = 2,300 ms; flip angle (FA) = 9°; image matrix = 240 mm × 256 mm; slice thickness = 1 mm; 208 slices. Then, whole-brain functional images were acquired using a T2*-weighted echo planar imaging (EPI) sequence sensitive to BOLD contrast with the following specifications: TE = 30 ms; TR = 800 ms; FA = 52°; image matrix = 90 mm x 90 mm; slice thickness = 2.4 mm; slice gap = 2.4 mm; 660 slices. Stimuli were presented with the Psychophysics Toolbox (Brainard, 1997) for MATLAB using an LCD AVOTEC projector onto a screen located behind the participant’s head and viewed through an integrated head-coil mirror.

Data preprocessing was carried out using the Configurable Pipeline for the Analysis of Connectomes (C-PAC version 1.6.0, https://fcp-indi.github.io/), an open-source software for the automated preprocessing and analysis of fMRI data (Craddock et al., 2013). C-PAC is implemented in Python using the *Nipype* pipelining library (Gorgolewski et al., 2011) that integrates tools from AFNI (Cox, 1996), FSL (Smith et al., 2004), and ANTs (Avants et al., 2008) to achieve high-throughput processing on high-performance computing systems. Head motion was assessed with an algorithm that quantifies the root mean square of the relative framewise displacement (FDRMS) between each volume of functional data (Smith et al., 2004). No participants were excluded for excessive head motion, which was defined *a priori* as mean FDRMS greater than 0.15 mm.

Anatomical images were resampled to RPI orientation and a non-linear transform between skull-on images and a 2mm MNI brain-only template was calculated using ANTs (Avants et al., 2008). Images were skull-stripped using AFNI’s 3dSkullStrip (Cox, 1996) and segmented into white matter (WM), grey matter (GM), and cerebrospinal fluid (CSF) using FSL’s FAST tool (Zhang et al., 2001). The resulting WM mask was multiplied by a WM prior map that was transformed into individual space using the inverse of the linear transforms previously calculated during the ANTs procedure. A CSF mask was multiplied by a ventricle map derived from the Harvard-Oxford atlas distributed with FSL (Makris et al., 2006). Skull-stripped images and grey matter tissue maps were written into MNI space at 2mm resolution.

Functional preprocessing began with resampling the data to RPI orientation and slice timing correction. Next, motion correction was performed using a two-stage approach in which the images were first coregistered to the mean fMRI and then a new mean was calculated and used as the target for a second coregistration (AFNI 3dvolreg; Cox & Jesmanowicz, 1999). A seven degree of freedom linear transform between the mean fMRI and the anatomical image was calculated using FSL’s implementation of boundary-based registration (Greve & Fischl, 2009). Nuisance variable regression (NVR) was performed on motion-corrected data using a second order polynomial, a 24-regressor model of motion (Friston et al., 1996), nuisance signals obtained from white matter (CompCor; Behzadi et al., 2007), and mean CSF signal. WM and CSF signals were extracted using the previously described masks after transforming the fMRI data to match them in 2mm space using the inverse of the linear fMRI-sMRI transform. The NVR procedure was performed with the inclusion of global signal as a nuisance regressor. The residuals of the NVR procedure were written into MNI space at 2.4mm resolution and subsequently smoothed using a 5mm FWHM kernel.

For all social attention functional scans, parcellation was performed by taking the framewise average of the voxelwise signals in each of 268 nodes from the Shen atlas (Shen et al., 2013). The Shen atlas is a functional parcellation that covers the whole brain, including cortex, subcortex, and cerebellum. For the Attend Faces and the Attend Houses conditions separately, we calculated z-scored Pearson correlation coefficients between the activity timecourses of all possible pairs of nodes to construct 268 x 268 symmetric functional connectivity matrices. Only the upper triangle of each functional connectivity matrix was extracted and vectorized, and the resulting 35,778 connections or edges served as input features to CPM.

### Connectome-based predictive modeling

We performed CPM using a 10-fold cross-validation procedure where the data was divided into 10 folds and models were trained using 9 of the 10 folds (Shen et al., 2017; Finn & Bandettini, 2021). The resulting models were tested on data from the held-out fold to generate predicted behavioral scores for all 6 or 7 participants in that fold. By iterating through all 10 folds, predicted behavioral scores were generated for all participants in the sample. This procedure was repeated for 100 iterations to assess sensitivity of model accuracy to different fold splits. Also, at each fold in the 10-fold cross-validation procedure, we residualized the target behavioral variable with respect to potential confound variables. The target behavioral variable was modeled as a linear combination of the potential confound variables, and the residuals of the model served as input features to CPM.

First, mass univariate correlation between the strength of each edge and the target behavioral score was performed in the training set. A threshold of |*r*| > 0.2 was applied so only the most significantly correlated edges were selected for further analysis. Previous research has shown that this threshold provides good accuracy with relatively sparse features (Finn et al., 2015). Then, the selected edges were divided into a positive and a negative tail based on the sign of their correlation with behavior. For each participant, the strength across all edges was summed into a single value, separately for the edges positively and negatively correlated with behavior.

Next, linear models were fit with network strength as the independent variable and the behavioral measure as the dependent variable. Lastly, positive and negative network strength for each participant in the test set were calculated by applying the masks defined by the training set to their functional connectivity data. Network strength served as input to the linear model to generate predicted behavioral scores for all participants in the test set.

Prediction accuracy of the models was measured as the Spearman correlation between observed or true behavior and predicted or model-generated behavior. Spearman correlation was used to assess prediction accuracy since successful rank prediction across participants was our goal. To assess the statistical significance of prediction accuracies, we generated a null distribution of expected accuracies due to chance by shuffling behavioral scores with respect to connectivity matrices and reperforming the entire analysis pipeline for a total of 1,000 randomizations. Then, we calculated a non-parametric p-value, which counts the number of times prediction accuracy for each of the 1,000 iterations of the null distribution exceeds the median prediction accuracy of the 100 true iterations.

For significant CPM models, we performed additional steps to interpret the functional anatomy of predictive networks. First, we averaged the number of times an edge was selected within each 10-fold run. Then, we averaged those fractions across all 100 iterations of 10-fold cross-validation. We limited visualization to edges that appeared significant in at least 97 percent of all the models to ensure we were considering only edges that were most robustly predictive of behavior. Trends that distinguished between positive and negative networks were also visualized by comparing relative numbers of edges between pairs of anatomically-defined macroscale regions.

## Results

### Prediction of social attention task performance during selective attention to faces

In our sample, d’ during the Attend Faces condition ranged from −0.56 to 2.80 (*M* = 1.53, *SD* = 0.72), demonstrating that this task is capable of generating significant variability in performance that is ideal for studying individual differences in social attention ability.

Additionally, d’ was not significantly correlated with age (*r* = −0.05, *p* = 0.693) and did not significantly differ between males (*M* = 1.38, *SD* = 0.84) and females (*M* = 1.61, *SD* = 0.63; *t* = - 1.18, *p* = 0.243). We focused analyses on the Attend Faces condition because we were specifically interested in functional connectivity during selective attention to faces and its relevance to autistic and socially anxious traits.

We first demonstrated that whole-brain functional connectivity patterns during the Attend Faces condition of the social attention task predicted d’ during the Attend Faces condition with significant accuracy. For this CPM model, observed and predicted d’ values were significantly correlated (median *r* = 0.40, *p* = 0.005; Fig. 1a). We included FDRMS as a covariate during CPM to control for the impact of head motion on d’ (*r* = −0.30, *p* = 0.013). The high-social attention network consisted of 882 edges whose strength predicted higher d’ across participants, and the low-social attention network consisted of 833 edges whose strength predicted lower d’ across participants (Fig. 1c). The twelve most highly-connected nodes in the high- and low-social attention networks were located in the occipital lobe, parietal lobe, temporal lobe, cerebellum, prefrontal cortex, motor strip, subcortex, and limbic regions (Tables 1a and 1b).

**Figure 1.**
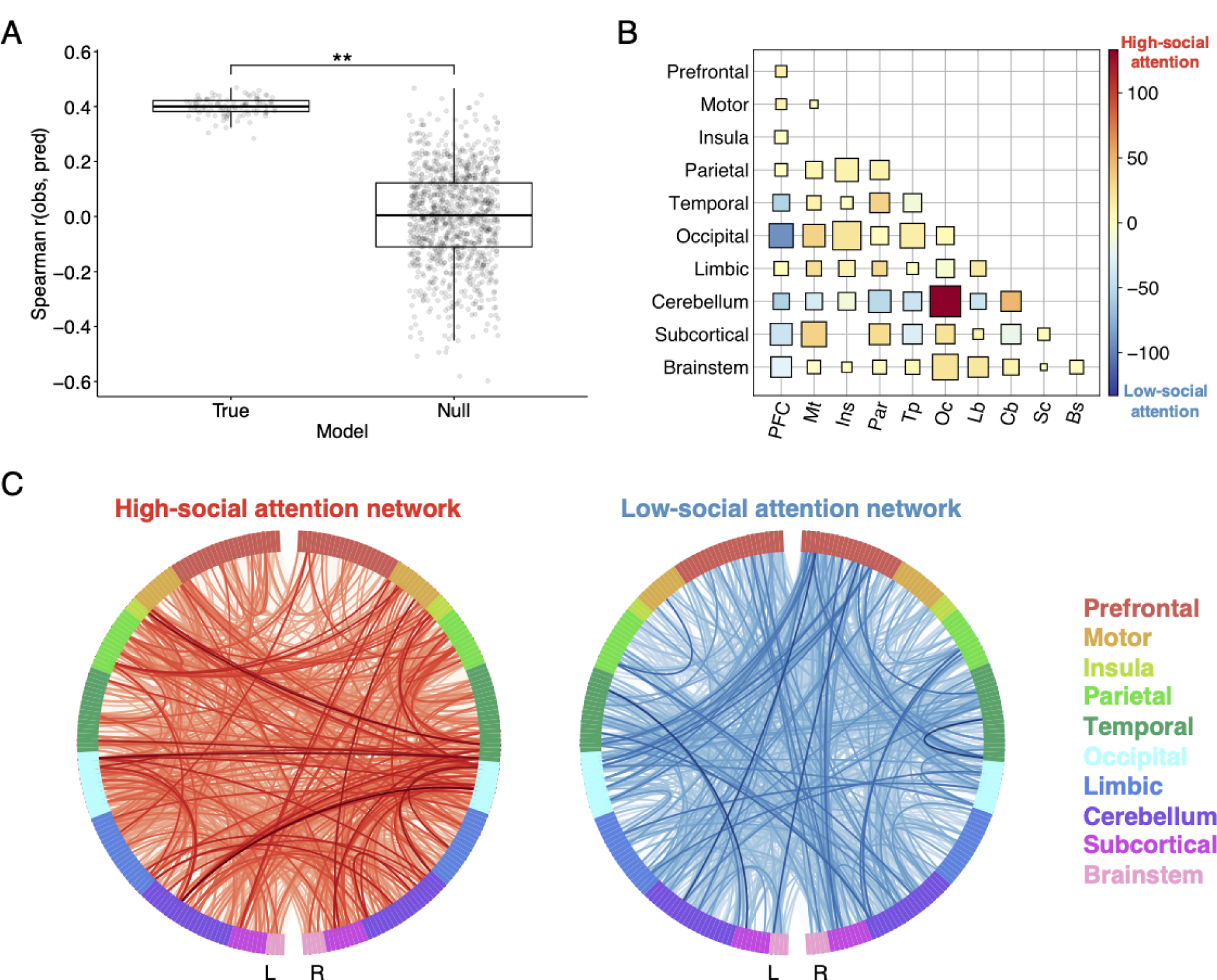
Connectome-based predictive model trained to predict d’ from functional connectivity during selective attention to faces. A) Prediction accuracy was measured as the Spearman correlation between observed and predicted scores (*y* axis). Results from 100 true iterations of 10-fold cross-validation are shown on the left and results from 1,000 permutations comprising the null distribution are shown on the right (*x* axis). Statistical significance was calculated by comparing the median prediction accuracy of the true models (horizontal line) to the null distribution; **p < 0.01. B) Lobewise visualization of edges in the high- and low-social attention networks. Size corresponds to the sum of edges in the high- and low-social attention networks standardized by the number of possible edges between each pair of regions. Color corresponds to the difference between edges in the high- and low-social attention networks, such that red corresponds to edges mostly in the high-social attention network and blue corresponds to edges mostly in the low-social attention network. C) The 882 edges in the high-social attention network (predicting higher d’ values) are visualized in red. The 833 edges in the low-social attention network (predicting lower d’ values) are visualized in blue. Darker lines correspond to edges with higher strength (absolute value).

**Table 1a.** High-social attention network key nodes. Key nodes are defined as those with the highest degree, or number of connections with other nodes. *K*, degree; Hem: L, left hemisphere, R, right hemisphere; BA, Brodmann area; MNI, Montreal Neurological Institute coordinates.

| Node # | Node name | *K* | Hem | Lobe | Network | BA | MNI |
| --- | --- | --- | --- | --- | --- | --- | --- |
| 208 | VisualAssoc | 37 | L | Occipital | Visual I | 19 | -16.79, -84.92, 33.04 |
| 42 | VisMotor | 33 | R | Parietal | Visual I | 7 | 14.82, -68.41, 34.88 |
| 212 | SecVisual | 32 | L | Occipital | Visual II | 18 | -10.83, -98.14, 7.63 |
| 213 | SecVisual | 28 | L | Occipital | Visual II | 18 | -14.72, -83.97, -13.06 |
| 49 | AngGyrus | 28 | R | Parietal | Default Mode | 39 | 41.39, -75.34, 27.98 |
| 203 | VisualAssoc | 26 | L | Occipital | Default Mode | 19 | -41.25, -75.44, 22.76 |
| 110 | Cerebellum | 22 | R | Cerebellum | Limbic | n/a | 21.07, -54.82, -23.76 |
| 206 | VisualAssoc | 20 | L | Occipital | Visual Association | 19 | -43.24, -70.37, -13.83 |
| 25 | PreMot+SuppMot | 20 | R | MotorStrip | Motor | 6 | 6.97, -8.07, 52.92 |
| 222 | VentPostCing | 19 | L | Limbic | Default Mode | 23 | -8.6, -58.79, 17.61 |
| 180 | SupramargGyr | 19 | L | Parietal | Motor | 40 | -42.21, -31.23, 15.89 |
| 71 | Fusiform | 19 | R | Temporal | Visual Association | 37 | 41.65, -45.73, -22.64 |

**Table 1b.** Low-social attention network key nodes. Key nodes are defined as those with the highest degree, or number of connections with other nodes. *K*, degree; Hem: L, left hemisphere, R, right hemisphere; BA, Brodmann area; MNI, Montreal Neurological Institute coordinates.

| Node # | Node name | *K* | Hem | Lobe | Network | BA | MNI |
| --- | --- | --- | --- | --- | --- | --- | --- |
| 206 | VisualAssoc | 28 | L | Occipital | Visual Association | 19 | -43.24, -70.37, -13.83 |
| 71 | Fusiform | 25 | R | Temporal | Visual Association | 37 | 41.65, -45.73, -22.64 |
| 139 | AntPFC | 22 | L | Prefrontal | Fronto-parietal | 10 | -18.21, 56.99, -14.27 |
| 110 | Cerebellum | 22 | R | Cerebellum | Limbic | n/a | 21.07, -54.82, -23.76 |
| 149 | FrontEyeFields | 21 | L | Prefrontal | Medial Frontal | 8 | -39.35, 17.2, 46.7 |
| 102 | Cerebellum | 21 | R | Cerebellum | Visual II | n/a | 39.09, -74.91, -29.7 |
| 261 | Putamen | 20 | L | Subcortical | Basal Ganglia | n/a | -24.78, 5.62, -0.08 |
| 253 | Cerebellum | 20 | L | Cerebellum | Cerebellum | n/a | -26.31, -69.52, -30.6 |
| 180 | SupramargGyr | 20 | L | Parietal | Motor | 40 | -42.21, -31.23, 15.89 |
| 152 | ParsOrbitalis | 20 | L | Prefrontal | Limbic | 47 | -28.35, 36.03, -15.64 |
| 143 | AntPFC | 20 | L | Prefrontal | Fronto-parietal | 10 | -42.72, 47.25, -6.93 |
| 138 | AntPFC | 20 | L | Prefrontal | Default Mode | 10 | -6.93, 48.31, -5.71 |

Additionally, we found that connections between the cerebellum and occipital cortex primarily predicted better social attention task performance since there were 139 of those connections in the high-social attention network and only 6 of those connections in the low-social network (Fig. 1b). Meanwhile, connections between prefrontal cortex and occipital cortex predicted worse social attention task performance as there were 95 of those connections in the low-social attention network and only 3 of those connections in the high-social attention network (Fig. 1b).

### Estimation of three latent autistic and socially anxious trait communities

We estimated three distinct communities from bootEGA that correspond to social skills, behavioral inflexibility, and social anxiety domains (Fig. 2). High scores on the social skills community were associated with less interest and enjoyment in social interaction and less effective communication with others, while high scores on the behavioral inflexibility community were associated with difficulty adapting to change and switching attentional focus. Lastly, high scores on the social anxiety community were associated with more fear of social situations and of scrutiny from others. Eigenvalue-eigenvector decomposition corroborates the three-community dimensionality assessment from bootEGA, and the three latent variable communities were stably estimated such that items consistently loaded onto the same factor across repeated iterations. Also, the three latent variable communities were not orthogonal since items from different communities had non-zero weights onto other communities. We did not expect any of these communities to be fully independent of each other, as the social skills and behavioral inflexibility communities are both aspects of the autistic phenotype and there is high comorbidity between autistic and socially anxious phenotypes (Maddox & White, 2015). We did not find behavioral evidence of a relationship between social attention abilities and subclinical autistic and socially anxious traits, since d’ during the Attend Faces condition of the social attention task was not significantly correlated with social skills (*r* = −0.07, *p* = 0.581), behavioral inflexibility (*r* = 0.001, *p* = 0.993), or social anxiety (*r* = 0.01, *p* = 0.908).

**Figure 2.**
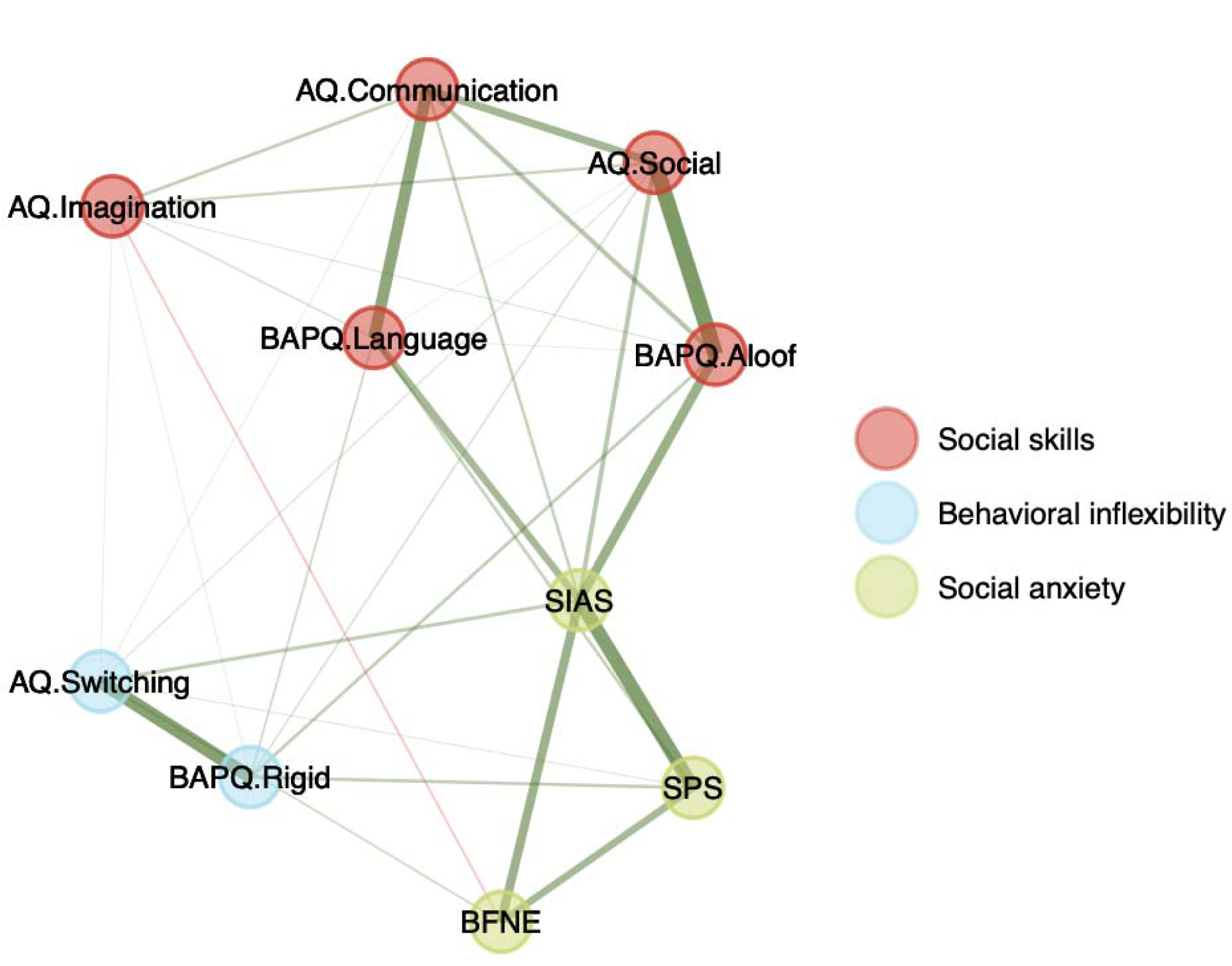
Three latent communities estimated from autistic and socially anxious trait questionnaire data using bootstrap exploratory graph analysis. Distinct color circles reflect distinct communities. Positive edges are visualized in green and negative edges are visualized in red. Thicker lines correspond to edges with higher strength (absolute value).

### Prediction of social skills from the high-social attention network

Next, we utilized functional connectivity between brain regions predictive of better social attention task performance to further examine the association between social attention and behavioral phenotypes. Specifically, CPM was performed using functional connectivity strength of the 882 edges from the high-social attention network to predict social skills, behavioral inflexibility, and social anxiety scores. We demonstrated that functional connectivity patterns between the 882 edges from the high-social attention network predicted social skills with significant accuracy, as observed and predicted social skills scores were significantly correlated (median *r* = 0.35, *p* = 0.023; Fig. 3a). This CPM model included behavioral inflexibility scores (*r* = 0.83, *p* < .001) and social anxiety scores (*r* = 0.86, *p* < .001) as covariates. The low-social skills network consisted of 9 edges whose strength predicted more autistic-like social skills across participants, and the high-social skills network consisted of 15 edges whose strength predicted less autistic-like social skills across participants (Fig. 3c). We found that connections between occipital cortex and motor strip primarily predicted fewer social skills since there were 5 of those connections in the low-social skills network and only 2 of those connections in the high-social skills network (Fig. 3b). Meanwhile, we found that connections between temporal cortex and parietal cortex primarily predicted more social skills since there were 4 of those connections in the high-social skills network and none of those connections in the low-social skills network (Fig. 3b). Additionally, the high-social skills network exhibited right hemisphere lateralization, as 11 of the 15 connections were within the right hemisphere.

**Figure 3.**
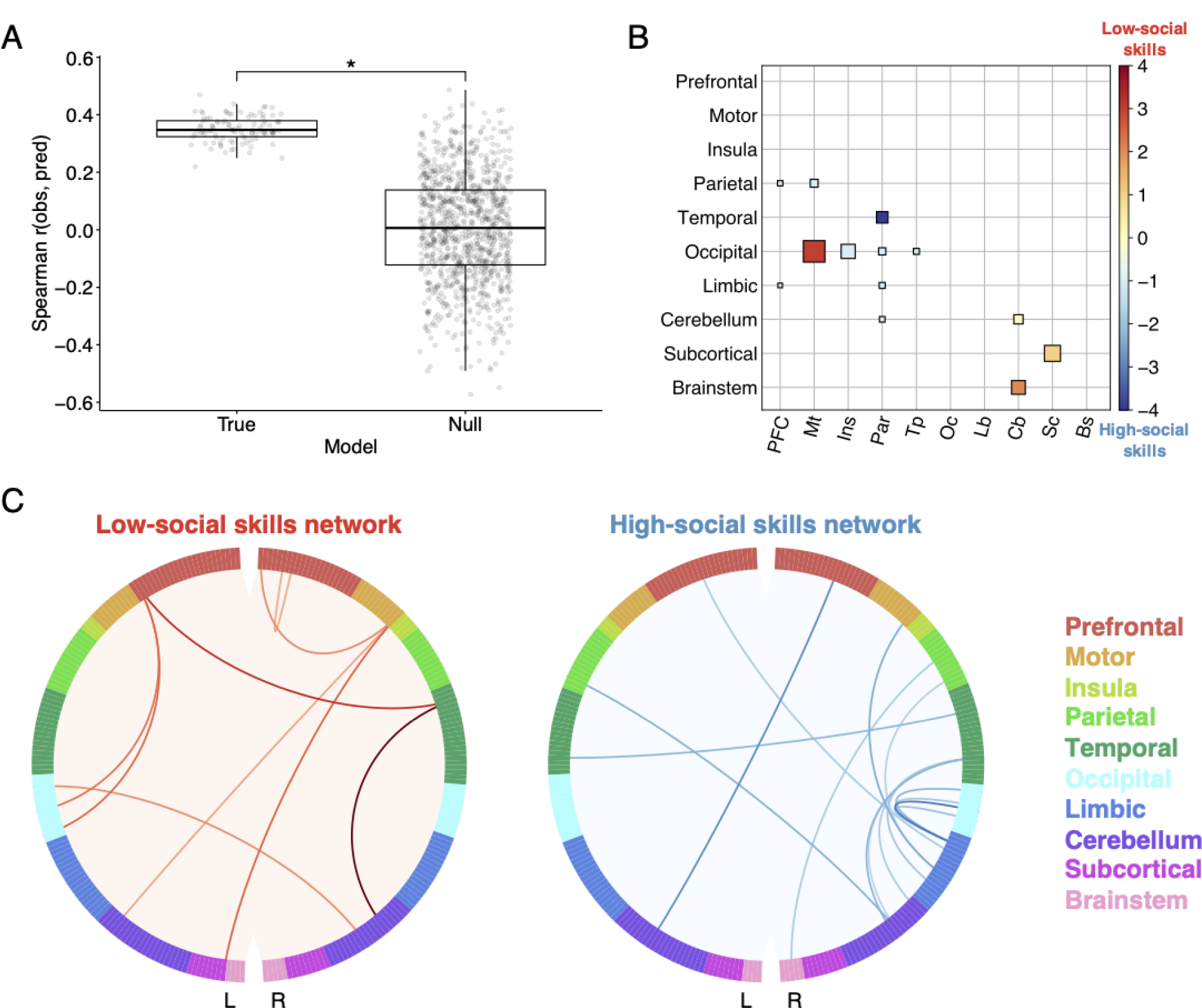
Connectome-based predictive model trained to predict social skills from functional connectivity within the high-social attention network. A) Prediction accuracy was measured as the Spearman correlation between observed and predicted scores (*y* axis). Results from 100 true iterations of 10-fold cross-validation are shown on the left and results from 1,000 permutations comprising the null distribution are shown on the right (*x* axis). Statistical significance was calculated by comparing the median prediction accuracy of the true models (horizontal line) to the null distribution; *p < 0.05. B) Lobewise visualization of edges in the low- and high-social skills networks. Size corresponds to the sum of edges in the low- and high-social skills networks standardized by the number of possible edges between each pair of regions. Color corresponds to the difference between edges in the low- and high-social skills networks, such that red corresponds to edges mostly in the low-social skills network and blue corresponds to edges mostly in the high-social skills network. C) The 9 edges in the low-social skills network (predicting more autistic-like social skills scores) are visualized in red. The 15 edges in the high-social skills network (predicting less autistic-like social skills scores) are visualized in blue. Darker lines correspond to edges with higher strength (absolute value).

### Prediction of behavioral inflexibility from the high-social attention network

We demonstrated that functional connectivity patterns between the 882 edges from the high-social attention network failed to predict behavioral inflexibility with significant accuracy, as observed and predicted behavioral inflexibility scores were not significantly correlated (median *r* = 0.15, *p* = 0.212). This CPM model included social skills scores and social anxiety scores (*r* = 0.81, *p* < .001) as covariates.

### Prediction of social anxiety from the high-social attention network

We demonstrated that functional connectivity patterns between the 882 edges from the high-social attention network failed to predict social anxiety with significant accuracy, as observed and predicted social anxiety scores were not significantly correlated (median *r* = −0.12, *p* = 0.760). This CPM model included social skills scores and behavioral inflexibility scores as covariates.

## Discussion

We identified and quantified functional brain networks predictive of individual differences in social skill in a neurotypical sample. First, we utilized whole-brain functional connectivity during selective attention to faces to accurately predict performance on an fMRI social attention task. Then, the significant edges identified in step one were used to predict scores on three latent communities of autistic and socially anxious traits. Latent community structure was derived from an EGA model constructed using self-report questionnaire data completed by a large independent sample. We found that connectivity strength within the social attention connectome predicted social skills, but not behavioral inflexibility or social anxiety. These results have implications for our basic understanding of the neural systems underlying individual differences in social attention abilities in the general population, which are related to varying levels of social skill.

While the functional anatomy of predictive edges uncovered in this study was widespread, a few notable connections were important for distinguishing individuals on the basis of their social attention abilities. Specifically, connectivity between the cerebellum and the occipital cortex, which is involved in the processing of visual information during attention (Posner & Petersen, 1990), was most predictive of better performance on the social attention task. Although the cerebellum is traditionally implicated in the prediction and execution of motor sequences, recent evidence indicates that the cerebellum also supports numerous social cognitive processes through adaptive prediction (Van Overwalle et al., 2014; Frosch, Mittal & D’Mello, 2022). Our findings suggest that individuals with more cerebellar-occipital connectivity performed better on the social attention task, perhaps indicating that they were more efficient at updating prior information to process rapidly changing social information in the context of visually distracting information. Meanwhile, connectivity between the prefrontal cortex and the occipital cortex was most predictive of worse performance on the social attention task. DLPFC is a prefrontal region involved in effortful processing and attentional control (Hopfinger et al., 2000), and previous research found that additional recruitment of DLPFC during selective attention to faces allowed both autistic and neurotypical individuals to compensate for intrinsically reduced salience of social information (Herrington et al., 2015; Puglia et al., 2018). The current results extend these prior findings by demonstrating that connectivity with the broader prefrontal cortex, not just DLPFC, is predictive of social attention ability in neurotypical individuals with different degrees of subclinical autistic traits.

Additionally, we were able to identify functional connections that predicted social skill, but not behavioral inflexibility or social anxiety. Here, we reduced our search space from connections distributed throughout the whole brain to those connections that significantly predicted performance on our experimental task, and then identified a subset of those connections as significant predictors of individual variability in self-reported social skills.

Connectivity between the motor strip and the occipital cortex was most predictive of lower social skills and more difficulty with social communication and interaction. Meanwhile, connectivity between the temporal lobe and the parietal lobe, as well as right hemisphere lateralization, was most predictive of more social skills and less difficulty with social communication and interaction. The right temporoparietal junction (TPJ), a brain region where the temporal and parietal lobes intersect in the right hemisphere, plays a critical role in both low- and high-level aspects of social cognition (Saxe & Kanwisher, 2003; Decety & Lamm, 2007). For example, the right TPJ is involved in reorienting attention to salient stimuli, as well as in theory of mind and empathy.

One potential limitation of this study is the relatively small sample size for research on individual differences. A recent paper by Marek and colleagues suggests that thousands of participants are required to reliably replicate associations between individual differences in brain function and complex phenotypes (Marek et al., 2022). However, by leveraging a challenging functional task and appropriate behavioral measures, our approach allowed CPM to uncover associations between individual differences in whole-brain functional connectivity patterns and social skill. The social attention fMRI task used in this study contributed to successful behavioral predictions by amplifying relevant variability in whole-brain signal (Greene et al., 2018). The one-back selective social attention task employed here was used previously to successfully distinguish between the neural responses of autistic and neurotypical individuals (Herrington et al., 2015), as well between a group of neurotypical individuals on the basis of task performance (Puglia et al., 2018). Additionally, the identification of specific social trait and behavioral domains through rigorous phenotyping of participants contributed to successful behavioral predictions. Autism is a complex phenotype, and it is critical that research aims to understand how the wide range of heterogeneity manifests by characterizing subtypes of autism (Lombardo, Lai & Baron-Cohen, 2019). Therefore, we focused on the prediction of specific autistic trait domains instead of the entire autistic phenotype. With this approach, we provide evidence that functional brain networks associated with social attention task performance also carry meaningful information about social skills.

Future work will show that functional brain networks predictive of social attention ability and social skills identified in this sample can generalize to predict social behavior in independent samples. The present study excluded autistic individuals, therefore limiting our ability to discuss social capabilities across the full autistic trait continuum. Ideally, both neurotypical and clinical populations should be included to further capture variability in key social cognitive processes.

Though this type of data collection may prove challenging for individuals that struggle with the experimental task, advances in naturalistic viewing paradigms with high-valence social content present a solution because, similar to task-based fMRI data, functional connectivity measured during naturalistic viewing yields more accurate predictions of trait-like phenotypes compared to resting-state data (Finn & Bandettini, 2021). Therefore, the social attention connectome established here can be applied to naturalistic viewing fMRI data from a novel group of individuals with or without autism or other developmental disorders to make predictions about social skills. The social attention connectome established here can similarly be applied to examine how the emergence of these networks is related to social skills in a pediatric sample in which many social behaviors are still developing. Importantly, this research may identify individuals early in life with an increased risk for social difficulties.

In summary, we demonstrated that individual variability in whole-brain functional connectivity reliably predicted social attention abilities and that a subset of those functional connections extended to also predict social skills. The use of an fMRI social attention task that generated variability in behavioral performance combined with the identification of subtypes of autistic and socially anxious traits led to the success of brain-based predictive models. This research identified a potentially generalizable neuromarker of social attention ability that ultimately predicts social interaction and communication skills that are essential to the development of fulfilling, meaningful social relationships.

## Acknowledgements

This research was supported by the National Science Foundation Grant 1657726.

